# Vehicle-mounted cameras reveal negative impact of the Fukushima Daiichi nuclear power plant accident on large bird abundance via paddy field abandonment

**DOI:** 10.1101/2023.05.31.540128

**Authors:** Nao Kumada, Keita Fukasawa, Akira Yoshioka, Naoe Tsuda, Hirofumi Ouchi

**Affiliations:** Biodiversity Division, National Institute for Environmental Studies, Tsukuba, Ibaraki, Japan; Fukushima Regional Collaborative Research Center, National Institute for Environmental Studies, Miharu, Fukushima, Japan; Center for Climate Change Adaptation, National Institute for Environmental Studies, Tsukuba, Ibaraki, Japan

**Keywords:** Abandoned paddy fields, Vehicle-mounted video camera, Agricultural birds, Distance sampling, Roadside survey, UAV

## Abstract

Farmland bird populations are declining due to farmland abandonment and agricultural intensification. Effective conservation strategies require appropriate monitoring approaches, including efficient and scalable survey methods. In this study, we developed a large-scale monitoring method for herons and egrets (Ardeidae) using vehicle-mounted video cameras and distance sampling models that incorporate location uncertainty. The survey was conducted in and around the evacuation zone of the Fukushima Daiichi nuclear power plant accident. A total of 7,031 km of roadside video footage was recorded, covering 24.41 km² of farmland. Ardeidae were observed only outside the evacuation zone and were entirely absent within it. Predicted abundance differed greatly between areas inside (0.0279 ± 0.0307/km²) and outside (4.57 ± 5.36/km²) the evacuation zone. Incorporating location uncertainty into the distance sampling model had little effect on the estimates (4.57 ± 5.36 vs. 4.51 ± 5.29/km² with and without integrating location uncertainty, respectively). This suggests that our video-based roadside survey method is robust to location uncertainty in structured landscapes such as Japanese rice paddies. The accuracy may be attributed to the study system, where levees and roads divide paddy fields, limiting potential error in observer-target distances Our method can be applied to other open habitats, such as natural grasslands and wetlands, especially in areas lacking artificial markers, by incorporating measurement uncertainty into the model. This combination of roadside surveys with vehicle-mounted cameras and distance sampling provides a practical and transferable approach for monitoring large-bodied species in open landscapes, enhancing both the efficiency and spatial coverage of biodiversity assessments.

**Highlights:** Broad-scale farmland abandonment following the Fukushima Daiichi nuclear power plant accident led to sharp declines in large farmland bird abundance.

Efficient large-scale roadside surveys were conducted using vehicle-mounted video cameras.

Ardeidae abundance was estimated using distance sampling models that incorporated location uncertainty.

The survey method is robust and applicable to open habitats such as farmlands, grasslands, and wetlands.

## 1 Introduction

Population sizes of many farmland birds are declining due to agricultural intensification and the abandonment of cultivation (Fujioka, Don Lee, Kurechi, & Yoshida, 2010; Fuller et al., 1995; Siriwardena et al., 1998; Stanton, Morrissey, & Clark, 2018). To implement appropriate conservation strategies for farmland birds, information on their abundances and related environmental factors is essential. Various methods have been developed for monitoring farmland birds, including point count, transect count, territory mapping, and capture-recapture methods (Birrer et al., 2007; Chiron, Filippi-Codaccioni, Jiguet, & Devictor, 2010; Gillings & Fuller, 1998). These methods are often labor-intensive and costly to conduct over a large area. Cost-effective methods for monitoring are needed to better understand population patterns and environmental drivers of farmland birds at a large spatial scale and hence to develop effective conservation measures.

In Japan, the Fukushima Daiichi nuclear power plant accident in 2011 led to uninhabited areas due to an evacuation order for the contaminated area, and all agricultural activities within those areas were stopped (Yoshioka, Mishima, & Fukasawa, 2015). Although observable effects of ionizing radiation at the population level on terrestrial organisms were unlikely (UNSCARE, 2017), land abandonment in the evacuation zone can have indirect effects. These environmental changes are thought to have a significant impact on the organisms that use farmland. Therefore, extensive monitoring in the farmland throughout the region is needed to evaluate the effect of farmland abandonment in the evacuation zone. However, with respect to radiation protection, it is advisable for researchers not to stay in the evacuation zone for a long time, necessitating an efficient monitoring scheme. Main farmland in the region is paddy fields, which are used by a lot of herons and egrets (Ardeidae). Therefore, they are suitable indicator species of the paddy field environment. They are the main predators in paddy fields and are particularly vulnerable to the abandonment of paddy fields (Amano & Katayama, 2009; Lane & Fujioka, 1998). Their large size and conspicuous whitish body color make Ardeidae foraging in paddy fields easily detectable from roads.

Vehicle-mounted cameras offer a novel and efficient approach for wildlife monitoring along roads. These systems enable continuous, non-intrusive image collection over large areas while minimizing the need for prolonged field presence. The roadside survey approach, involving species observations from moving vehicles, has the capacity to collect data for species abundance indices across broad areas at low cost (Abella, Spencer, Hoines, & Nazarchyk, 2009; Catry, Moreira, Deus, Silva, & Águas, 2015; Milton & Dean, 1998). Recently, vehicle-mounted camera footage—including from platforms such as Google Street View—has been used to monitor birds and other organisms in various habitats (Deus, Silva, Catry, Rocha, & Moreira, 2016; Olea & Mateo-Tomás, 2013; Rousselet et al., 2013). However, applications for estimating animal density have been limited to areas with clearly defined and uninterrupted backgrounds (e.g., birds nesting on cliffs) , and challenges remain in open habitats such as farmlands, where detection ranges are uncertain. Ignoring this uncertainty can lead to biased estimates of abundance and environmental effects (Buckland, Marsden, & Green, 2008).

Distance sampling can resolve the issue of bias in abundance estimation by video-based roadside survey. Distance sampling estimates animal density or abundance from the distance between the observer and the detected individual, accounting for the decrease in detection probability due to distance (Buckland et al., 2001; Fox et al., 2017; Hara, Yamaji, Yokota, Thaitakoo, & Sampei, 2018; Howe, Buckland, Després-Einspenner, & Kühl, 2017; Kvasnes, Pedersen, & Nilsen, 2018; Stober, Prieto-Gonzalez, Smith, Marques, & Thomas, 2017). Distance sampling can be used to estimate trends in farmland bird populations in roadside surveys. However, there might be uncertainty in the distance between the observer and the detected individual measured from video images. For robust abundance estimation, an independent field test of location uncertainty in distance sampling is required (Hefley, Boyle, & Mohankumar, 2020).

Comparison of the observer–target distances measured by roadside survey and unmanned aerial vehicles (UAVs) is expected to be a feasible independent field test. UAVs have become an effective tool for monitoring animal populations and individual locations (Iwamoto, Nogami, Ichinose, & Takeda, 2022; Witczuk, Pagacz, Zmarz, & Cypel, 2018). Although their application to large spatial areas remains costly, this method is highly accurate because UAVs can measure animal positions from almost directly above. If the same individuals are observed by both roadside survey and UAV, location uncertainty in roadside survey can be quantified. Integrating location uncertainty obtained by these small-scale measurements into a stochastic distance sampling model will allow us to moderate cost–accuracy trade-offs and estimate abundance at a large spatial scale.

The purpose of this study was to estimate the population density of Ardeidae in rice fields in and around the evacuation zone of the Fukushima Daiichi nuclear power plant accident by roadside survey. We surveyed the locations of Ardeidae by using a vehicle-mounted camera and then estimated population densities by using a distance sampling model incorporating location errors. The magnitude of location errors by roadside survey was determined by a field test comparing roadside survey with UAV-mounted cameras. To evaluate the importance of accounting for location uncertainty, population densities were compared between analyses with and without location errors. We demonstrated the impact of paddy field abandonment due to the evacuation order by comparing the population density with a counter-factual scenario without an evacuation order.

## 2 Materials and methods

### 2.1 Roadside survey of Ardeidae around the evacuation zone

The roadside survey with vehicle-mounted cameras of abandoned and cultivated farmlands was conducted in eastern Fukushima, which includes the evacuation zone of the Fukushima Daiichi nuclear power plant accident (Fig. 1). The evacuation zone was designated after the Fukushima Daiichi nuclear power plant accident in March 2011 and is divided into three categories according to radiation levels: Difficult-to-Return Area, Restricted-Residential Area, and Preparing for the Lifting of the Evacuation Order Area (Cabinet Office, Government of Japan, 2013). In all three categories, cropland cultivation did not resume until 2016, except in some test fields. The study area covered farmlands within 18 municipalities located inside and outside the evacuation zone.

**Figure 1.**
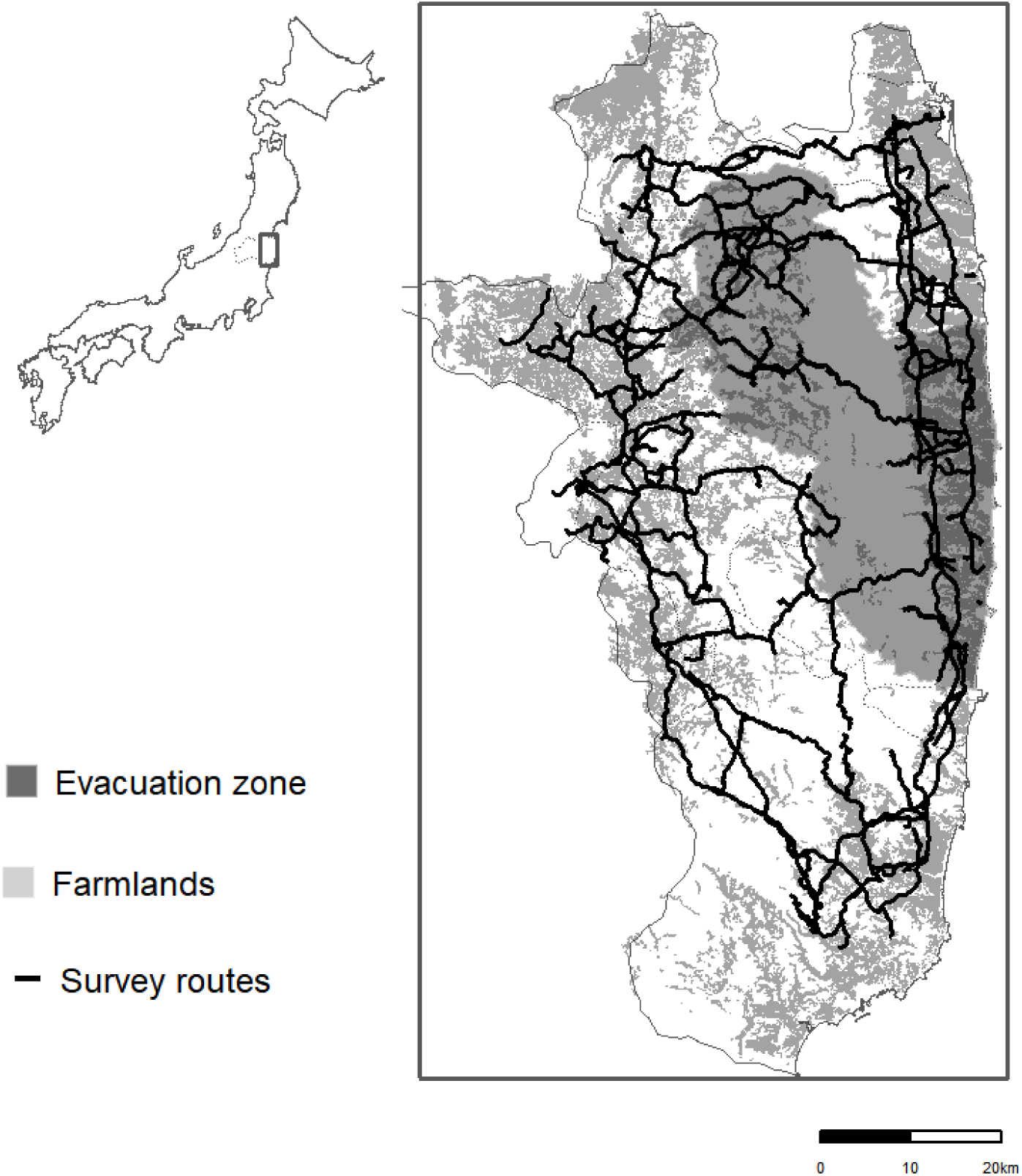
Map of survey routes (black lines) in Eastern Fukushima prefecture. Dark grey area shows the evacuation zone of the Fukushima Daiichi nuclear power plant accident. Light grey areas show farmland polygons.

From early May to early July between 2014 and 2016, roadside video recording surveys were conducted on farmlands by vehicle-mounted cameras. Video cameras with GPS (VIRB Elite or VIRB X; Garmin Ltd.) were mounted on the side windows of a vehicle, which was driven on roads adjacent to farmlands for video recording (Fig. 2). Videos recorded by the two camera models (VIRB Elite or VIRB X) simultaneously at the same location showed no obvious differences. Although the video cameras were mounted on both sides of a car in many cases, some surveys were carried out using a video camera mounted on one side only. The GPS trajectories of the car were also recorded simultaneously. Videos were shot at 1920 × 1080 p/30 fps for all models and saved in mp4 file format. The speed of the car during the survey varied depending on the speed limit (60 km/h or less) and road congestion; however, the frame rate was sufficiently high to detect Ardeidae under all speeds in the study.

**Figure 2.**
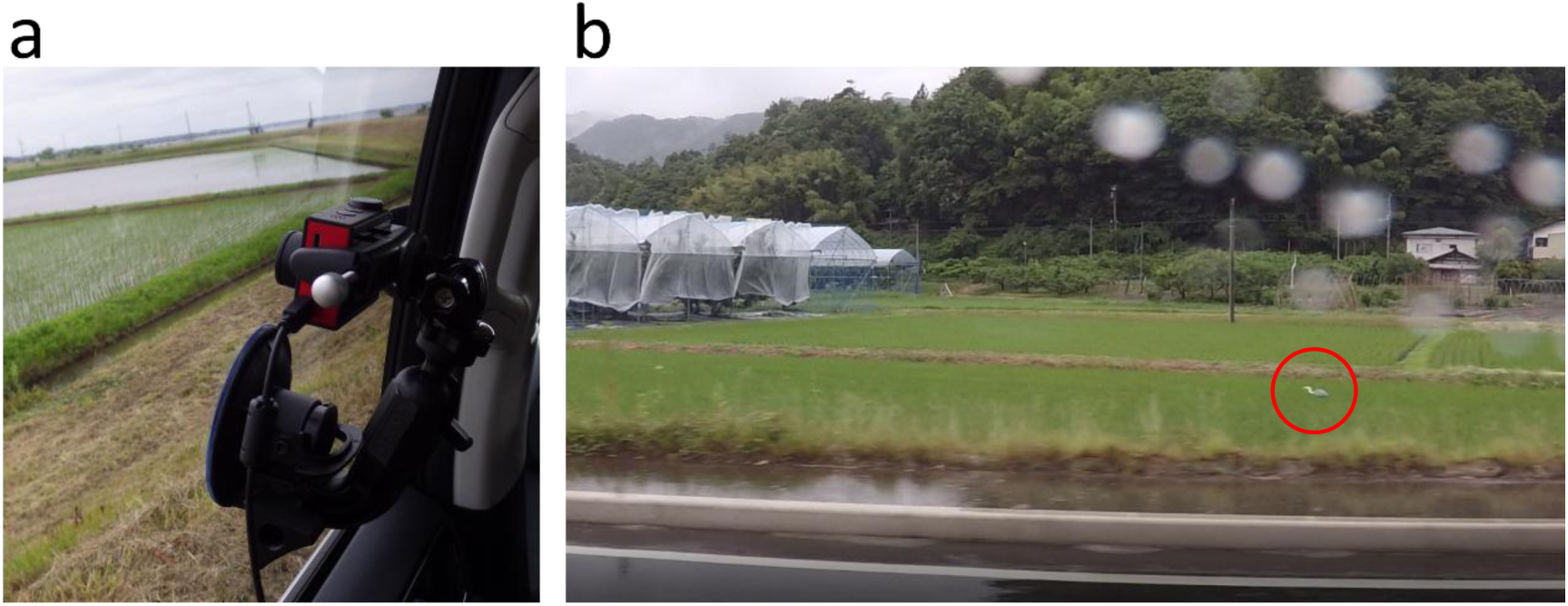
Photo showing a video camera with GPS mounted on a side window of a vehicle (a) and photo, captured from a video recorded by a vehicle-mounted camera, showing Ardeidae (in a red circle) in a paddy field (b).

To identify farmland within the viewshed from the vehicle-mounted cameras, the farmland polygons in the direction of the camera and not hidden by obstacles such as forest and houses were selected as follows (Fig. 3). Farmland polygons in this area were created from aerial photographs taken in 2006–2015 (see Appendix 1 for detailed methods). The farmland polygons included upland fields, pastures, and abandoned farmlands, and more than 70% of the farmland area was paddy fields in Fukushima prefecture before the nuclear power plant accident (in 2010) (Ministry of Agriculture, Forestry and Fisheries, 2020). First, we extracted the farmland polygons in the direction of the camera and intersecting within a straight-line distance of 10 m from the trajectories of the car; these were defined as the viewshed. Ten meters was set as an approximate threshold for adjacency (i.e., no obstacles) between the road and farmland. Second, we sequentially included farmlands that were within 10 meters from the viewshed obtained in the previous step. Farmlands were not included in the viewshed when the distance from the movement trajectory was more than 500 m because this was too far to detect Ardeidae. We visually verified that the selection criteria could effectively extract farmlands within the viewshed of the camera. For subsequent analysis, only paddy field polygons were used, which were selected from the identified polygons based on existing information (Ministry of Agriculture, Forestry and Fisheries, 2021) and aerial photographs.

**Figure 3.**
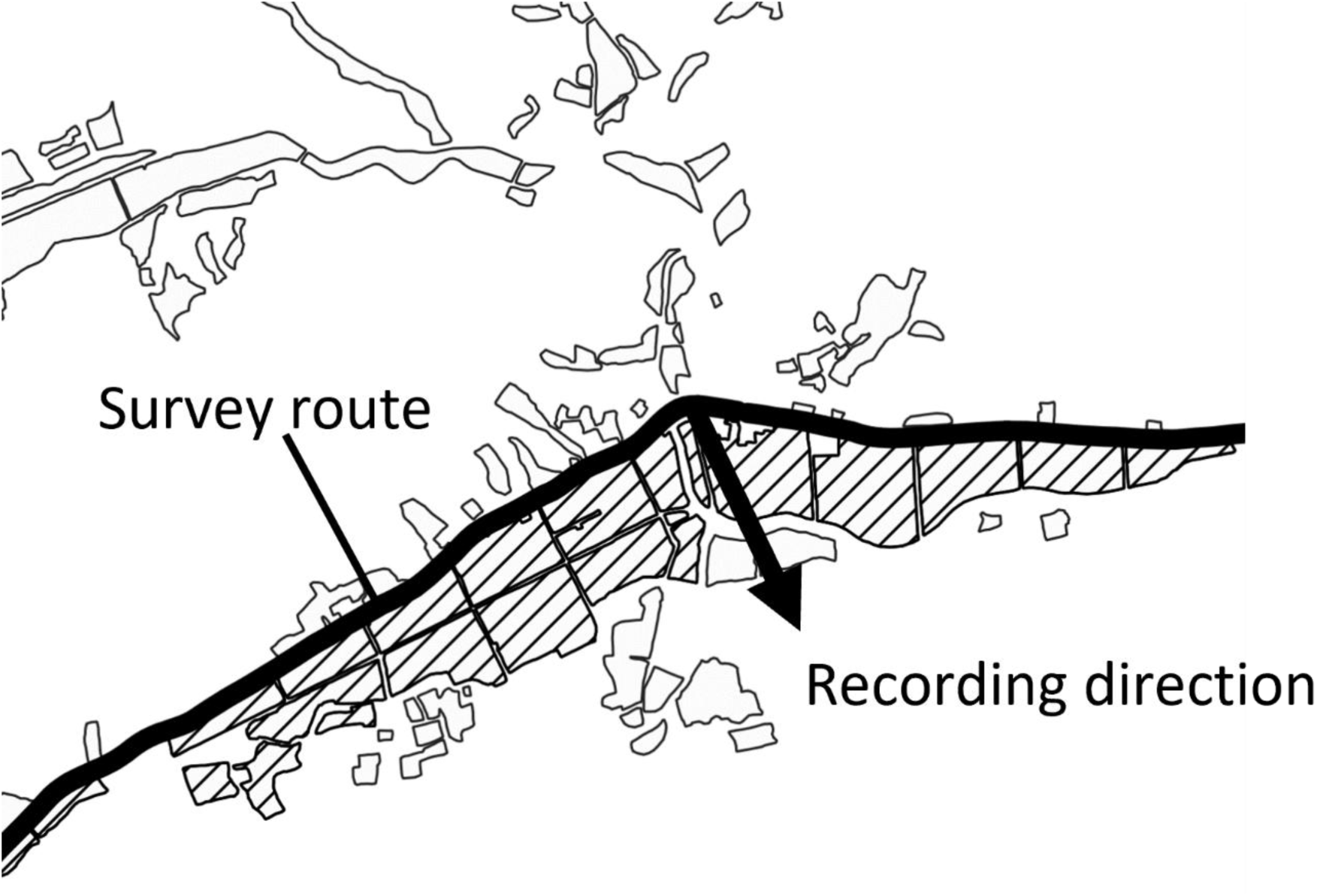
Illustration of the selection of farmlands within the viewshed. A survey route (bold line) and farmland polygons (surrounding lines) are shown. The polygons that meet the criteria (see text) are selected as viewsheds (shaded areas).

We played the videos back and searched for Ardeidae in paddy fields. Then, we determined the locations on the map following the same method used to test location errors described above. Because it was difficult to identify species from the video images correctly, except for grey heron, we merged all species of Ardeidae for further analyses. The following Ardeidae may be found in the study area: grey heron *Ardea cinerea*, great egret *A. cinerea*, intermediate egret *A. intermedia*, little egret *Egretta garzetta*, and cattle egret *Bubulcus ibis*. All of these Ardeidae are colonial and breed in the summer. In addition, flooding status (i.e., flooded or not) was recorded for the paddy fields where Ardeidae were found.

### 2.2 Field test of location errors of Ardeidae using UAVs

Location errors of the vehicle-mounted camera were validated by comparison with the accurate values determined by UAV images (UAV-truth). The survey was conducted on paddy fields in two regions, the district of Inashiki in Ibaraki Prefecture and the city of Minamisoma in Fukushima Prefecture. These paddy fields are about 5 square kilometers and are used for foraging by Ardeidae. The survey was conducted in Inashiki in July 2019 and May 2020 and in Minamisoma in June 2020. During the survey period, Ardeidae in the paddy fields were recorded simultaneously by a vehicle-mounted video camera (VIRB X; Garmin Ltd.) and a video camera in a UAV (Phantom 4; DJI). The GPS trajectories of vehicle-mounted video cameras were recorded by VIRB X simultaneously.

Locations of Ardeidae were determined from the movies captured by the vehicle-mounted cameras. If the researcher observed Ardeidae in the video, the movement trajectory information recorded at the time the video was taken was used to determine the location of the camera. Then, using information on the camera location with orthographical aerial imagery layers as background maps (GSI Tiles) and views surrounding Ardeidae (paddy levees, houses, telegraph poles, etc.) in the video image, the locations of Ardeidae were manually determined and recorded using QGIS ver.3.4.3.

To obtain correct locations of Ardeidae, georeferenced images obtained by the UAV were used. Georeferencing was performed using Georeferencer (Transformation type: Projective, Resampling method: Linear) in QGIS. The measurement errors of locations obtained by vehicle-mounted cameras were calculated as the Euclidean distances between the locations of the Ardeidae determined by the vehicle-mounted cameras and the UAV-truth values.

### 2.3 Statistical analysis

We employed a fully model-based approach to estimate the abundance of Ardeidae and the effect of land abandonment based on a distance sampling considering individual location uncertainty. As our observations consisted of the camera-based distance sampling results with location uncertainty and the independent field test of location errors based on correct locations determined by UAV, we integrated the likelihoods and estimated all parameters simultaneously.

A common approach for model-based distance sampling without location uncertainty is to treat the occurrence of individuals in relation to the distance from an observer as a realization of an inhomogeneous Poisson process (IPP) (Johnson, Laake, & Ver Hoef, 2010). The likelihood of distance sampling without location uncertainty is expressed by the following equation (Johnson, Laake, & Ver Hoef, 2010).

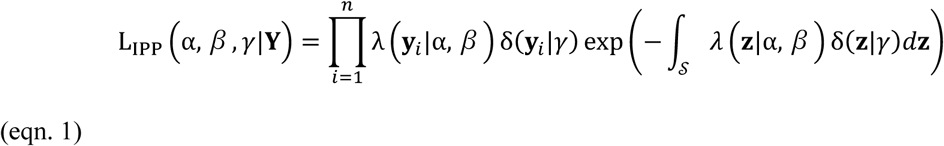

where **Y** = (**y**_1_, **y**_2_, …, **y***_n_*) are true locations of observed individuals, (*α,* **β**, *γ*) are model parameters, and 𝒮 is the spatial domain(s) that is potentially observable. λ(**z**|*α*, *β*) is an intensity function of the population density at location **z** and is commonly specified as follows.

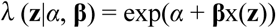

where α is the intercept, **β** = (*β*_1_,…,*β_p_*)′ is a vector of regression coefficients, and x(**z**) is a vector of covariates on population density at location **z**. δ(**z**|*γ*) is the detection probability, which is expressed as a thinning function as follows.

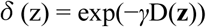

where D(**z**) is the Euclidian distance from an observer to location **z**.

Let us consider the case when observed location **Y**ʹ = (**y**ʹ_1_, **y**ʹ_2_, …, **y**ʹ*_n_*) is assumed to vary around the true location **Y** due to location error. The likelihood of IPP incorporating location uncertainty can be obtained by integrating out **y***_i_* over the surveyed region (Hefley et al., 2020).

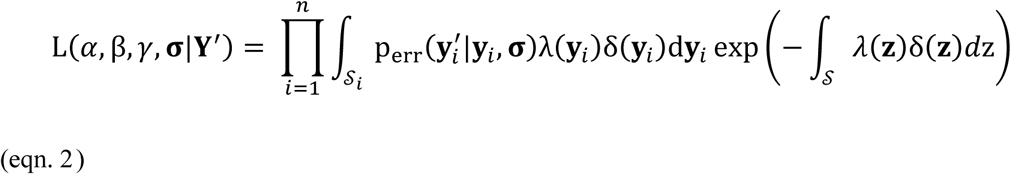

parameters *α,* **β**, *γ* were not included on the right side for simplicity) where p_err_(**y**ʹ|**y**, **σ**) is a probability distribution of location error, **σ** is a vector of location error parameters, and 𝒮_i_ is subset of 𝒮 in which **y**_i_ exist. For simplicity, we assume that the location error is isotropic and normally distributed.

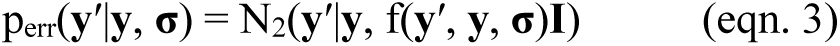

where f(**y**ʹ, **y**, **σ**) is a function of the standard deviation depending on the true location. In general, the greater the distance from the observer, the lower the accuracy of the observation position. The relationship was assumed to be described by a power function of Euclidean distances between **y**ʹ and **y**, D(**y**ʹ, **y**), with two parameters **σ** = (*σ*_0_, *σ*_1_) as follows.

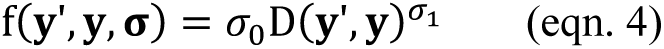

The likelihood function required integration over spatial domains, which is difficult to solve analytically. A practical approach is discretizing spatial domains into grid cells and replacing integration with piecewise summation as follows.

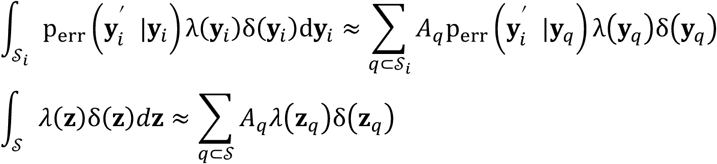

where **z***_q_* and *A_q_* are the centroid and area of grid cell *q*, respectively.

The location error parameter **σ** in the likelihood function described above is not identifiable without an independent survey of location error. In this study, the location error and UAV-truth were included simultaneously in the distance sampling model for the estimation of all parameters. When the true location is available, the error distribution (eqn. 3 and 4) can be written equivalently by the Rayleigh distribution (i.e., Weibull distribution with shape parameter 2) for the Euclidean distance between the observed and true location.

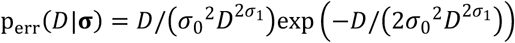

Given the vector of distances obtained by the field test D(**y**ʹ, **y**) = (D(*y*’_1_, *y*_1_), …, *D*(*y’_m_*, *y_m_*)), the joint likelihood is given as follows.

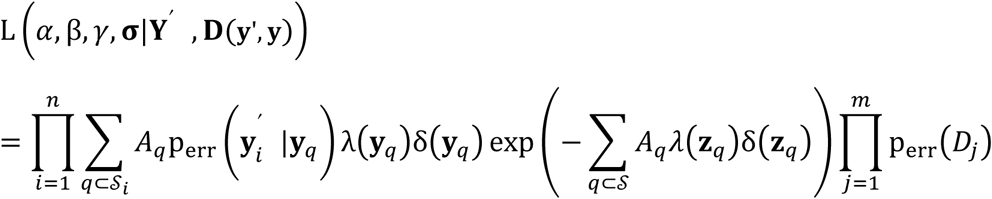

Both the maximum likelihood method and Bayesian method can be used for parameter estimation. When the Bayesian method is applied, prior information and constraints on parameters can be incorporated as a prior distribution.

We estimated the effects of the evacuation order (evacuation zone or not) (Cabinet Office, Government of Japan, 2013), the area of water bodies (rivers and lakes) (Ogawa et al., 2013) around the farmlands, and the year of the study on the abundance of Ardeidae. We defined farmland polygons within the viewshed of the camera as a multi-part domain 𝒮 and discretized it into grid cells with a resolution of 10 m. Then, these variables and the Euclidean distances from the camera were assigned to each cell. Even when location error is taken into account, the error is reasonably assumed to be within a paddy field polygon because paddy levees act as sharp borders dividing paddy field polygons. Therefore, we assumed that the subdomain 𝒮*_i_* containing the true location of the observed individual *i* was restricted to a paddy field polygon where the individual was observed.

The IPP model is often limited by overfitting, and it is common to apply regression shrinkage techniques. In this study, we applied the Bayesian default prior (Gelman, Jakulin, Pittau, & Su, 2007) to stabilize the coefficients and to prevent overfitting. We applied a weakly informative prior, Student’s *t*-test (df = 5, mean = 0, scale = 2.5), to regression coefficients **β** to avoid overfitting. For other parameters, improper flat priors p(*θ*) = 1 were used. The maximum *a posteriori* (MAP) estimate of each parameter was calculated using a quasi-Newton algorithm.

To estimate credible intervals of parameters, we obtained posterior samples by importance sampling (Gelman et al., 2013) with a proposal distribution that approximates the posterior. For the proposal distribution, we used a multivariate *t*-distribution with four degrees of freedom whose mean vector and scale matrix are the MAP estimate and the inverse of the Hessian matrix at the MAP, respectively. We drew 10,000 samples from the proposal distribution and obtained 3,000 posterior samples.

We calculated the abundance of Ardeidae by applying MAP estimates of each parameter to grid cells in paddy field polygons throughout the study area, with and without accounting for measurement error, and compared the two abundance results. The predicted values were aggregated for each 1-km square grid. To evaluate the effect of paddy field abandonment in the evacuation zone on the population size of Ardeidae, we predicted the population size in the study area under a counter-factual scenario in which no evacuation order had been given. All analyses in this study were implemented using R (R Core Team, 2021) (Appendix 2).

## 3 Results

### 3.1 Field test of location error of Ardeidae

In total, 78 Ardeidae (24 in July 2019, 31 in May 2020 in Inashiki and 23 in June 2020 in Minamisoma) were located with GIS by the vehicle-mounted camera and the camera on the UAV. The Euclidean distance between the locations of the Ardeidae determined by the vehicle-mounted camera and the true locations determined by the UAV was defined as the measurement error (Fig. 4). The location errors were increased with the distance between the observer and the Ardeidae.

**Figure 4.**
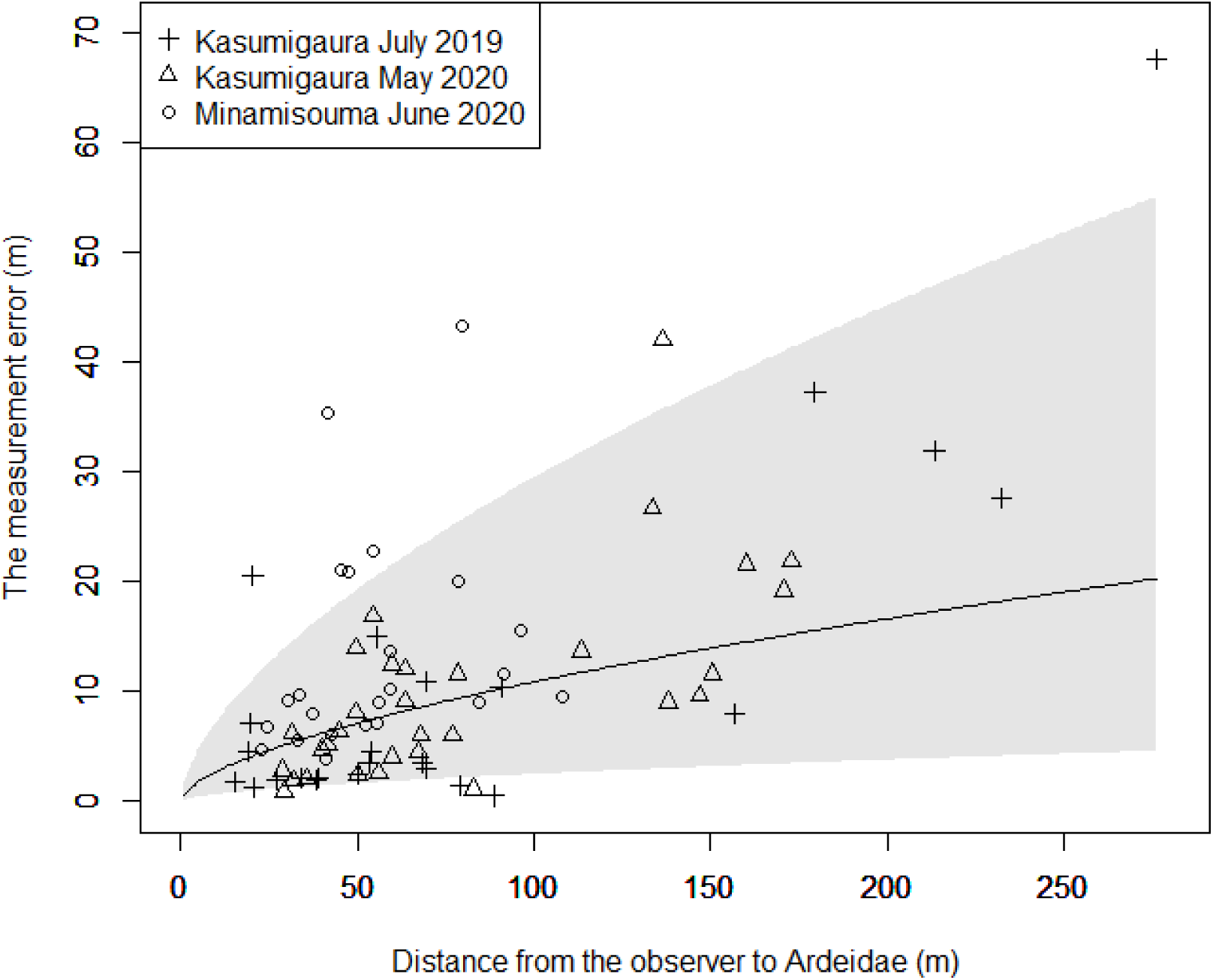
Relationship between the distance from cameras to Ardeidae and location errors. Black line and grey area show the maximum likelihood estimates and 95% credible interval for the regression function.

### 3.2 Roadside survey with vehicle-mounted cameras and abundance estimates of Ardeidae

The total distances traveled while recording with cameras were 3,675 km in the evacuation zone and 3,356 km outside the evacuation zone. The areas of farmland polygons (including abandoned fields in response to the evacuation order), selected as the viewshed, were 10.07 km^2^ in and 14.34 km^2^ around the evacuation zone.

A total of 56 individual Ardeidae were observed around the evacuation zone. No Ardeidae were observed in the evacuation zone (Fig. 5). All Ardeidae were found in flooded paddy fields. The frequency distribution of observed distances to Ardeidae was more concentrated near zero than the distribution of distances to roads for all grid cells of paddy field polygons. Although all polygons had a median distance of 133.3 m from roads, the polygons containing Ardeidae discovery sites had a median distance of 47.1 m and a maximum of 104.4 m (Fig. 6).

**Figure 5.**
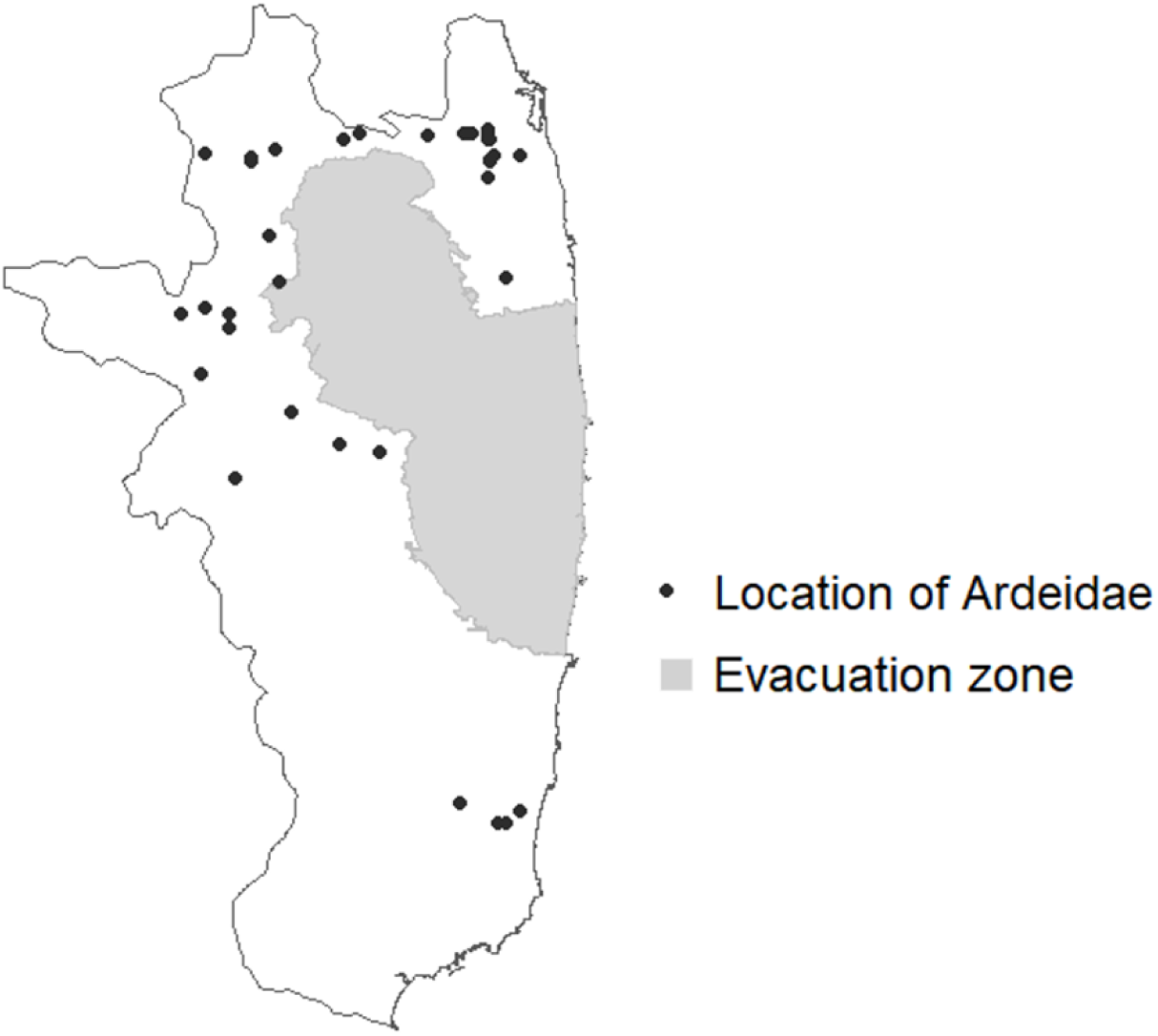
Location of Ardeidae observed by roadside survey.

**Figure 6.**
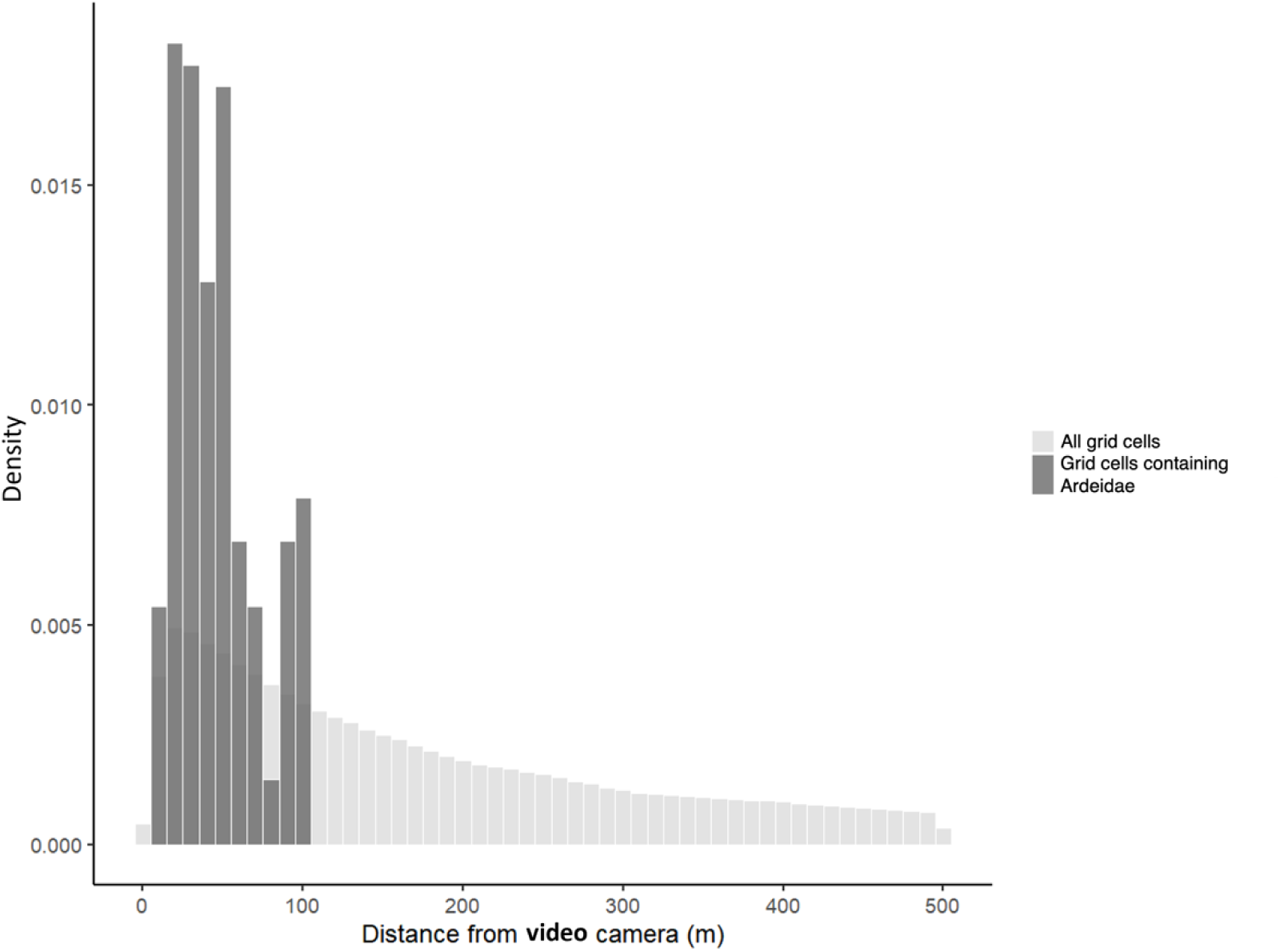
Histograms of the distance from the video camera to all grid cells of farmland polygons (light grey bars) and grid cells containing Ardeidae (dark grey bars).

The MAP estimates and summary statistics obtained by importance sampling are shown in Table 1. The water body area had a positive association with the abundance of Ardeidae. Abundance was lower in the evacuation zone than outside the zone. Although the signs (i.e., positive or negative) of parameter estimates were similar between the models with and without the location error, the parameter values differed slightly. The distance from the camera had a negative association with the probability of detection (Fig. 4). The probability decreased to half at 35.4 m (95% credible interval: 29.6–42.8 m) from the camera. The scaling factor and power value when measurement errors are expressed as a power function of the distance from the camera were 0.644 and 0.614. The measurement errors tended to increase with increasing distance from the camera on the car but exhibited a concave downward trend.

**Table 1.** MAP estimates and summary statistics via importance sampling with and without the location error.

| Parameter | With the location error |  |  |  |  | Without the location error |  |  |  |  |
| --- | --- | --- | --- | --- | --- | --- | --- | --- | --- | --- |
|  | Map | Mean | S.D. | CI2.5 | CI97.5 | Map | Mean | S.D. | CI2.5 | CI97.5 |
| Intercept | 6.55 | 7.11 | 1.48 | 4.79 | 10.4 | 6.54 | 7.12 | 1.48 | 4.76 | 10.4 |
| Distance | -0.0190 | -0.0191 | 0.00180 | -0.0228 | -0.0156 | -0.0188 | -0.0189 | 0.00183 | -0.0227 | -0.0155 |
| Water | 0.567 | 0.561 | 0.0820 | 0.391 | 0.717 | 0.567 | 0.561 | 0.0823 | 0.392 | 0.716 |
| Evac zone | -5.28 | -5.86 | 1.46 | -9.02 | -3.59 | -5.28 | -5.89 | 1.45 | -9.14 | -3.60 |
| Year(2015) | 0.190 | 0.203 | 0.263 | -0.292 | 0.760 | 0.192 | 0.218 | 0.265 | -0.294 | 0.749 |
| Year(2016) | -0.876 | -0.863 | 0.270 | -1.37 | -0.316 | -0.875 | -0.853 | 0.267 | -1.36 | -0.325 |

The predicted abundance of Ardeidae were substantially larger outside the evacuation zone than inside the zone (Table 2, Fig. 7). This trend did not differ between the models with and without location error. The predicted total population of Ardeidae in the study area was 7,820 individuals. Assuming that the evacuation zone was never designated, the predicted population would have increased by 2,647 individuals to 10,467 individuals (Fig. 8).

**Figure 7.**
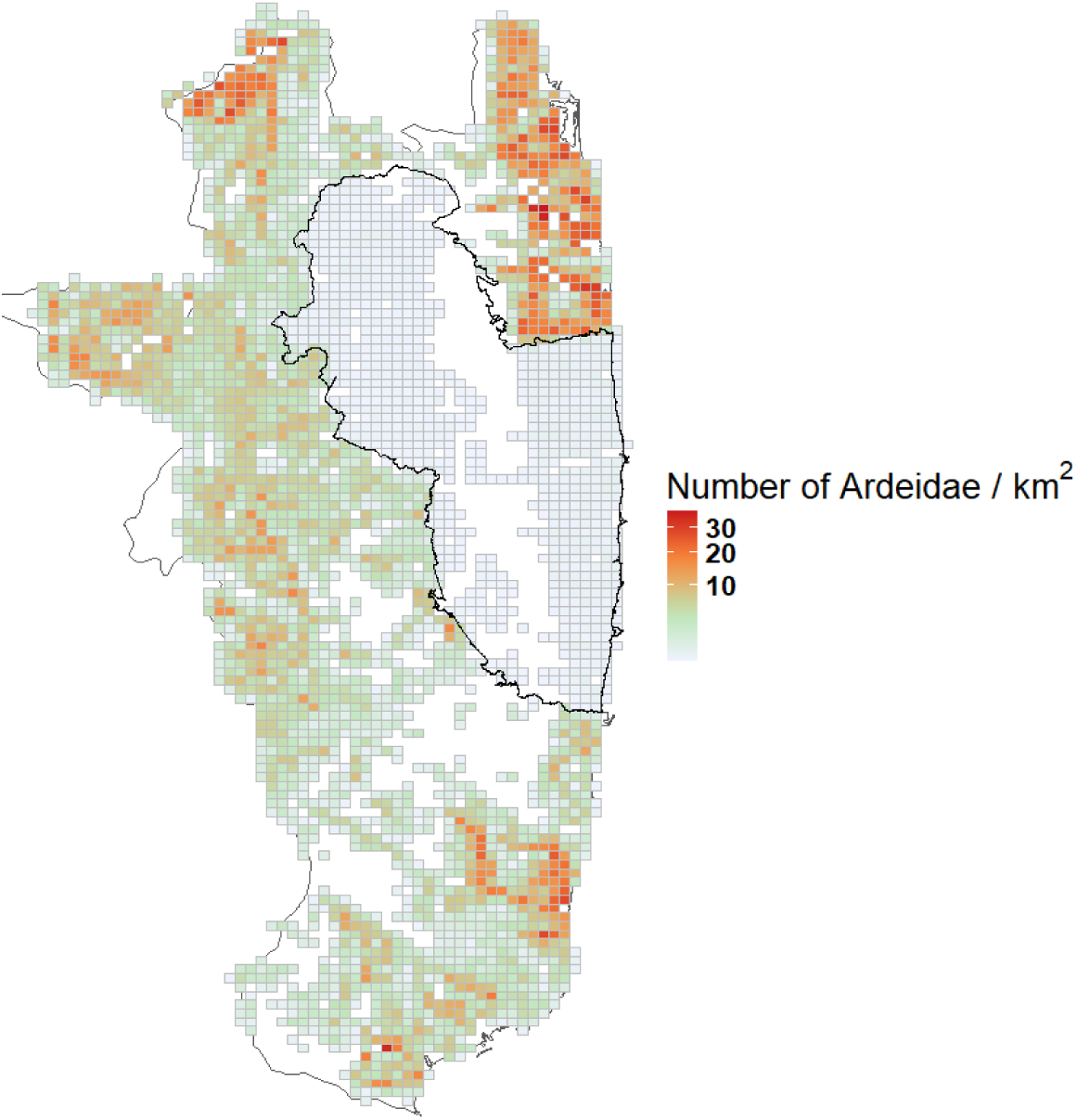
Prediction map of number of Ardeidae per square kilometer. Only grids that contain farmland area are included. The evacuation zone is circled by solid black lines.

**Figure 8.**
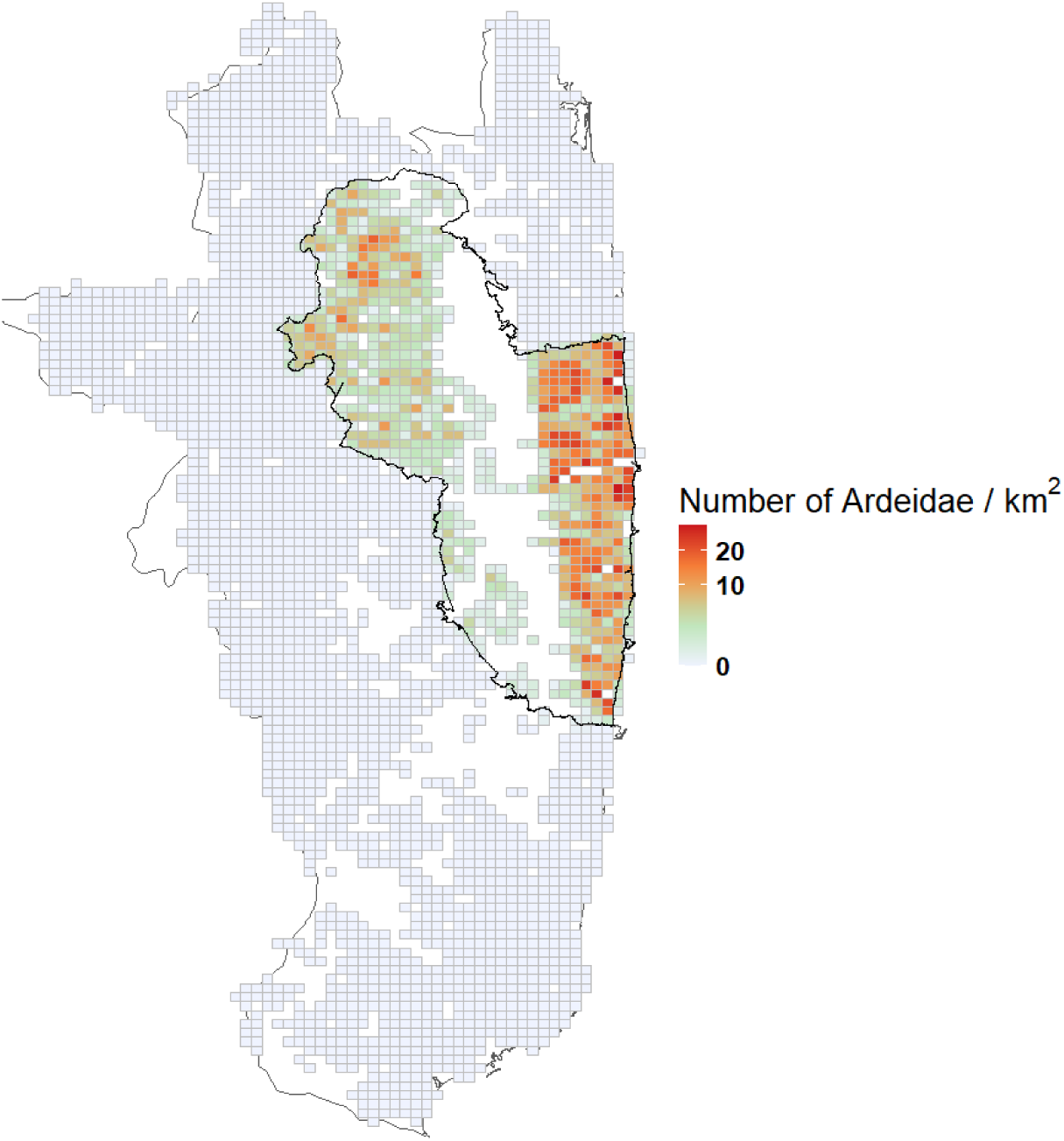
Map of the difference between the two predicted values for the numbers of Ardeidae per square kilometer. The actual value and the value obtained when assuming a counterfactual condition that the evacuation zone was not established were compared.

**Table 2.** Predicted values for the number of Ardeidae per square kilometer using the model with and without location error.

| Area category | With the location error |  | Without the location error |  |
| --- | --- | --- | --- | --- |
|  | Mean | S.D. | Mean | S.D. |
| Outside the evacuation zone | 4.57 | 5.36 | 4.51 | 5.29 |
| Inside the evacuation zone | 0.0279 | 0.0307 | 0.0276 | 0.0303 |

## 4 Discussion

We developed an efficient method for monitoring large birds in open areas by combining roadside surveys using vehicle-mounted cameras and distance sampling. By incorporating measurement error into the distance sampling framework, we addressed the challenge posed by the lack of a clearly defined observation area a common limitation of roadside surveys. The spatially explicit approach enabled us to estimate the impact of large-scale land abandonment on Ardeidae populations. Our survey revealed a large gap in Ardeidae distribution in the evacuation zone and allowed us to quantify the magnitude of this impact as a loss of abundance (Fig. 8, Table 2). The estimated number of foraging individuals around the evacuation zone was consistent with a previous field study that reported approximately 30 individuals per km^2^ (Lane & Fujioka, 1998) , supporting the reliability of our abundance estimates.

The direct effects of radiation exposure from the Fukushima Daiichi nuclear power plant accident on wildlife populations and ecosystems are not large enough to be observable at the population level (UNSCEARE 2017). However, the evacuation orders in response to the nuclear accident have resulted in short-term changes in land use and human activities (Tamaoki, 2016; Yoshioka et al., 2015), and these changes may have important impacts on wildlife populations and ecosystems (Fujimoto, Mitsunaga, & Takeuchi, 2015; Lyons, Okuda, Hamilton, Hinton, & Beasley, 2020; Matsushima, Ihara, Inaba, & Horiguchi, 2021; Yoshioka et al., 2015). The strong negative impact of the evacuation order on Ardeidae populations can be attributed to the decrease in food availability due to the widespread abandonment of paddy fields. Paddy field abandonment leads to the growth of dense vegetation, which is not a common foraging environment for Ardeidae (Katayama, Osawa, Amano, & Kusumoto, 2015; Maeda, 2005). In addition, Ardeidae feed on fish and frogs in flooded paddy fields (Tojo, 1996), which can be maintained by cultivation (Maeda, 2001). Our results clearly demonstrated that paddy field abandonment in evacuation zones affects the distribution of birds over a wide area. These findings are consistent with the observation that the long-term abandonment of farmland reduces the biodiversity of farmlands (Koshida & Katayama, 2018). Since 2016, the evacuation zone has been reduced and agricultural cultivation has begun to resume. In the future, the effect of resumption should be tested by continued monitoring following the protocols developed in this study.

While the roadside survey approach benefits from the ability to efficiently survey a wide area, the use of vehicle-mounted cameras will further improve efficiency. In this study, 237 hours of video were recorded to search for Ardeidae. In a field survey, experts would be required for the full time; however, the roadside survey method with vehicle-mounted cameras does not require expertise in bird identification in the field. In addition, the work of sorting paddy fields without Ardeidae, accounting for the majority of each video, does not require expertise; instead, experts only need to check images with suspected Ardeidae. Crowdsourcing this sorting process can promote the participation of birders in remote areas who are interested in the survey. In the future, automated counting may be possible (Norouzzadeh et al., 2018).

An accurate distance between the observer and target is required for distance sampling to estimate density; accordingly, when the distance accuracy could not be guaranteed, it is necessary to account for measurement error in distance sampling (Hefley et al., 2020). The incorporation of measurement error in the location of Ardeidae did not significantly affect density estimates. This result showed that the distance accuracy of roadside survey with vehicle-mounted cameras in our study was sufficient for density estimation by distance sampling. The robust performance of the roadside survey can be explained by the study system. The Ardeidae were found in paddy fields, and the location error could not exceed the size of one rice field separated by paths or levees. However, use of a UAV to estimate location errors for incorporation into a distance sampling model may be an effective approach in situations where measurement errors are likely such as natural grasslands and wetlands, where artificial markers are absent. Roadside survey by vehicle-mounted cameras may be a feasible strategy for various environments by using this method.

The detection probability depends on the distance from the observer as well as various covariates, in which case multiple-covariate distance sampling is appropriate (Marques & Buckland, 2003; Thomas et al., 2010). In this study, we did not consider the evacuation zone as a covariate, despite the potential effect of this parameter on the detection probability, because no Ardeidae were observed in the evacuation zone. However, we believe the absence of Ardeidae in the evacuation zone reflects differences in true population densities resulting from the abandonment of rice field because dense vegetation is generally not suitable for foraging by these species (Katayama, Osawa, Amano, & Kusumoto, 2015; Maeda, 2005). These species were not found in the evacuation zone, even near roads in which detectability is less affected by the vegetation.

The escape behavior of birds during surveys might biases population estimates (Hutto & Young, 2003; Roberts & Schnell, 2006). At paddy fields bordering roads, Ardeidae may fly away to avoid the approach of vehicles carrying cameras. Ardeidae at a distance from the camera can therefore be overlooked, which can lead to biases in the detection function and population estimates. However, the videos obtained in this study did not reveal any Ardeidae in flight, which could be a response to an approaching vehicle. In fact, waterbirds are not substantially impacted by approaching vehicles (McLeod, Guay, Taysom, Robinson, & Weston, 2013). Similarly, in this study, the effect of approaching vehicles was considered negligible for Ardeidae. If this method is applied to broader taxa, the use of fisheye lenses or 360-degree cameras may be a viable option to capture individuals before they are affected by an observer.

In our survey, we estimated the number of Ardeidae using paddy fields; however, these estimates cannot be interpreted as total counts for the study area. The survey was conducted during the breeding season, and one member of each nesting pair always stays in the nest. In addition, because Ardeidae fly over a wide area in search of good foraging sites, it is thought that they frequently move in and out of the evacuation zone and travel to and from the study area. It is necessary to consider whether the number of Ardeidae obtained in this study is an indicator of the quality of the paddy field as a foraging site. However, we believe that the density of Ardeidae foraging in paddy fields is a viable indicator of biodiversity in agricultural land because it is a common parameter for assessing the quality of paddy fields as foraging sites (Amano & Katayama, 2009; Katayama et al., 2019; Lane & Fujioka, 1998).

Another limitation of our study is that we were unable to identify the Ardeidae at the species level. We considered the total abundance of Ardeidae. The five species in the study area have overlapping ecological niches; they forage in rice fields and often share colonies (Mashiko & Toquenaga, 2014). In addition, the number of Ardeidae is related to the abandonment of rice field and taxon diversity and can therefore be used as an indicator of species richness on paddy fields (Katayama, Mashiko, Koshida, & Yamaura, 2021; Katayama et al., 2019; Koshida & Katayama, 2018). In the future, technological improvements such as increased camera resolution may make it possible to identify species and evaluate responses.

## 5 Conclusion

The combination of roadside surveys using vehicle-mounted cameras and distance sampling provided a reliable and efficient method to estimate the abundance and distribution of large birds in open agricultural landscapes. This approach allowed us to quantify the strong negative impact of paddy field abandonment on Ardeidae populations in the evacuation zone following the Fukushima Daiichi nuclear power plant accident.

Thanks to the structured landscape of paddy fields, our system was robust to location measurement error; however, by incorporating measurement error into the model, it can also be applied to other open habitats such as wetlands and grasslands. The approach is scalable, cost-efficient, and compatible with emerging technologies such as UAVs, street-level imagery platforms (e.g., Mapillary https://www.mapillary.com), and machine learning-based species recognition (Norouzzadeh et al., 2021) and distance estimation (Johanns, Haucke, & Steinhage, 2022). It holds promise for wide-scale biodiversity monitoring and conservation planning in agricultural systems and beyond.

## DATA ACCESSIBILITY

Data will be uploaded to the Dryad Digital Repository after acceptance.

## Acknowledgements

We are grateful to Fukushima Robot Test Field for helping field work. Yoshio Mishima contributed to create the farmland polygon data but declined to join as a co-author. This study was partially supported by JSPS KAKENHI Grant number 21H03656.

## Authors’ contributions

Nao Kumada, Keita Fukasawa and Akira Yoshioka conceived the ideas and designed the methodology; All the authors collected the data; Nao Kumada and Keita Fukasawa analysed the data; Nao Kumada, Keita Fukasawa and Akira Yoshioka led the writing of the manuscript. All the authors reviewed and contributed critically to the draft and gave final approval for publication.

## Conflict of interest

The authors declare no conflict of interest.

## Notes

### Competing Interest Statement

The authors have declared no competing interest.

### Summary of Updates

we have corrected errors in the original dataset and reanalyzed the data accordingly. The updated analyses have resulted in minor/moderate/substantial changes to the results presented in the manuscript, including revised figures and statistical estimates. These changes are reflected in the updated Abstract, Results, and Discussion sections.

